# Cooperative regulation of coupled oncoprotein translation and stability in triple-negative breast cancer by EGFR and CDK12

**DOI:** 10.1101/2021.03.03.433762

**Authors:** Hazel X. Ang, Natalia Sutiman, Xinyue L. Deng, Luke C. Bartelt, Qiang Chen, Alejandro Barrera, Jiaxing Lin, Jeff Sheng, Ian C. McDowell, Timothy E. Reddy, Christopher V. Nicchitta, Kris C. Wood

**Author notes:** Lead contacts (Christopher V. Nicchitta, Kris C. Wood).

## Abstract

Evidence has long suggested that epidermal growth factor receptor (EGFR) may play a prominent role in triple-negative breast cancer (TNBC) pathogenesis, but clinical trials of EGFR inhibitors have yielded disappointing results. Using a candidate drug screen, we discovered that inhibition of CDK12 dramatically sensitizes diverse models of TNBC to EGFR blockade. Instead of functioning through CDK12’s well-established roles proximal to transcription, this combination therapy drives cell death through the 4E-BP1-dependent suppression of the translation and consequent stability of driver oncoproteins, including MYC. A genome-wide CRISPR/Cas9 screen identified the CCR4-NOT complex as a major determinant of sensitivity to the combination therapy whose loss renders 4E-BP1 unresponsive to drug-induced dephosphorylation, rescuing MYC translational suppression and stability. The central roles of CCR4-NOT and 4E-BP1 in response to the combination therapy were further underscored by the observation of CNOT1 loss and rescue of 4E-BP1 phosphorylation in TNBC cells that naturally evolved therapy resistance. Thus, pharmacological inhibition of CDK12 reveals a long proposed EGFR dependence in TNBC that functions through the cooperative regulation of translation-coupled oncoprotein stability.

## INTRODUCTION

Triple-negative breast cancer (TNBC) is an aggressive subtype of the disease that constitutes 15-20% of all breast cancers. TNBCs are clinically defined by their lack of expression of the three main targetable receptors in breast cancer – the estrogen (ER), progesterone (PR) and human epidermal growth receptor-2 (HER2) receptors. The genetic and molecular heterogeneity within this pooled disease subtype has made patient stratification and targeted treatment particularly challenging (Lehmann et al., 2011; Lehmann et al., 2016; Foulkes et al., 2010; Dent et al., 2007). While attempting to identify oncogenic drivers in TNBC, immunohistochemical and large-scale genomic studies have suggested that EGFR signaling may be frequently activated and associated with poor prognosis (Park et al., 2014; Cancer Genome Atlas Network, 2012; Hoadley et al., 2007; Nielsen et al., 2004). These findings have long positioned EGFR as an intriguing target in TNBC; however, efforts to target EGFR in unselected TNBC patients have yielded low response rates (Baselga et al., 2005; Fountzilas et al., 2005; Carey et al., 2012; Baselga et al., 2013; Corkery et al., 2009; Ali and Wendt, 2017, Cruz-Gordillo et al., 2020). This is indicative of possible intrinsic resistance and its underlying mechanisms as a major impediment to more widespread use of EGFR inhibitors in TNBC.

Broad dysregulation of gene expression is one of the hallmarks of cancer, including TNBC, pointing to the potential utility of targeted therapeutics that alter gene regulation (Hanahan and Weinberg, 2011; Wang et al., 2015). Indeed, the development of specific inhibitors targeting multiple key players in transcriptional regulation has provided opportunities for novel therapeutic interventions (Bradner et al., 2017; Augert and MacPherson, 2014; Christensen et al., 2014). Cyclin-dependent kinase (CDK) 12 is a versatile kinase well known for regulating transcriptional and post-transcriptional processes, and has also recently been shown to play a role in the regulation of cap-dependent translation (Liang et al., 2015; Bösken et al., 2014; Greifenberg et al. 2016; Dubbury et al., 2018; Choi et al., 2019; Chirackal Manavalan et al. 2020; Choi et al., 2020). THZ531, a first-in-class selective inhibitor of CDK12 and CDK13, has been reported to suppress expression of genes that support malignant progression and induce apoptosis in cell line models (Zhang et al, 2016; Zeng et al, 2018). This work, and our broadening understanding of CDK12’s regulation of DNA damage response pathway genes, have driven the recent development of CDK12 as both a biomarker and therapeutic target (Lui et al., 2018; Wu et al., 2018; Choi et al., 2019; Chou et al., 2020; Liu et al., 2020; Blazek et al., 2011; Iniguez et al, 2018; Krajewska et al., 2019; Quereda et al, 2019; Abida et al., 2020; Wang et al., 2020). However, the functional interactions between CDK12 and most major oncogenic signaling pathways have remained largely unexplored.

Motivated by the hypothesis that CDK12 may functionally interact with major oncogenic signaling pathways in TNBC, we performed a candidate drug screen to identify synergistic drug combinations between THZ531 and inhibitors targeting possible oncogenic disease drivers. This work led to the unexpected finding that intrinsic resistance to EGFR inhibition in TNBC, a longstanding and unexplained observation, is mediated by CDK12. Definition of the mechanism underlying the profound synergy between EGFR and CDK12 inhibitors revealed that the stability of driver oncoproteins in TNBC is subject to translation-coupled regulation by these kinases, thereby nominating a mechanistically distinct approach for targeting oncogenic dependencies in this important disease subtype.

## RESULTS

### CDK12 inhibition sensitizes TNBC cells to EGFR inhibition

As there is no known single, targetable oncogenic driver in TNBC, we designed a panel of inhibitors targeting an array of key molecular pathways that are frequently implicated in cancer cell proliferation, survival, differentiation, and apoptosis. We tested two TNBC cell lines with this panel of inhibitors in the presence versus absence of a low, sublethal dose of THZ531, a selective inhibitor of CDK12/13. Both TNBC lines were markedly sensitized to the EGFR inhibitors gefitinib and lapatinib, as reflected by 10-100-fold reductions in GI_50_ values in the presence of THZ531 (Figure 1A). Further studies showed consistent THZ531-mediated sensitization to EGFR inhibitors across each member of a panel of eight diverse TNBC cell lines, decreasing their GI_50_ values to the sub-micromolar range in each case (Figure 1B and 1C). The sensitization effect was specific to TNBC cell lines and not observed in luminal breast cancer cell lines (BT474, SK-BR-3), or the immortalized mammary epithelial line, MCF10A (Figure 1C). Long-term combined EGFR and CDK12/13 inhibition suppressed colony formation in multiple TNBC cell lines (Figure 1D). Using an established analytic tool and commonly referenced Bliss independence model, additional quantitative analyses of our drug combination screening data confirmed synergy between EGFR inhibitors and THZ531 in multiple TNBC cell lines (Figure S1A) (Ianevski et al, 2020; Bliss, 1939). To confirm that the effects of THZ531 are on-target, and to distinguish between the functional effects of inhibiting CDK12 versus CDK13, we first used a stereoisomer and a derivative of THZ531, each of which spare CDK12 and CDK13 (THZ531R and THZ532, respectively), demonstrating that these compounds lose all activity when combined with gefitinib (Figure S1B) (Zhang et al., 2016). Additionally, we observed that the sensitizing effect of THZ531 was lost in cells expressing a mutant version of CDK12 (CDK12^AS^) (Bartkowiak et al., 2019) that is not inhibited by the drug (Figure S1C), suggesting that CDK12, and not CDK13, is the major target of the drug in this context. Collectively, these findings demonstrate that EGFR is an oncogenic driver in TNBC, and that intrinsic resistance to EGFR inhibitors in TNBC can be mitigated by CDK12 inhibition.

**Figure 1.**
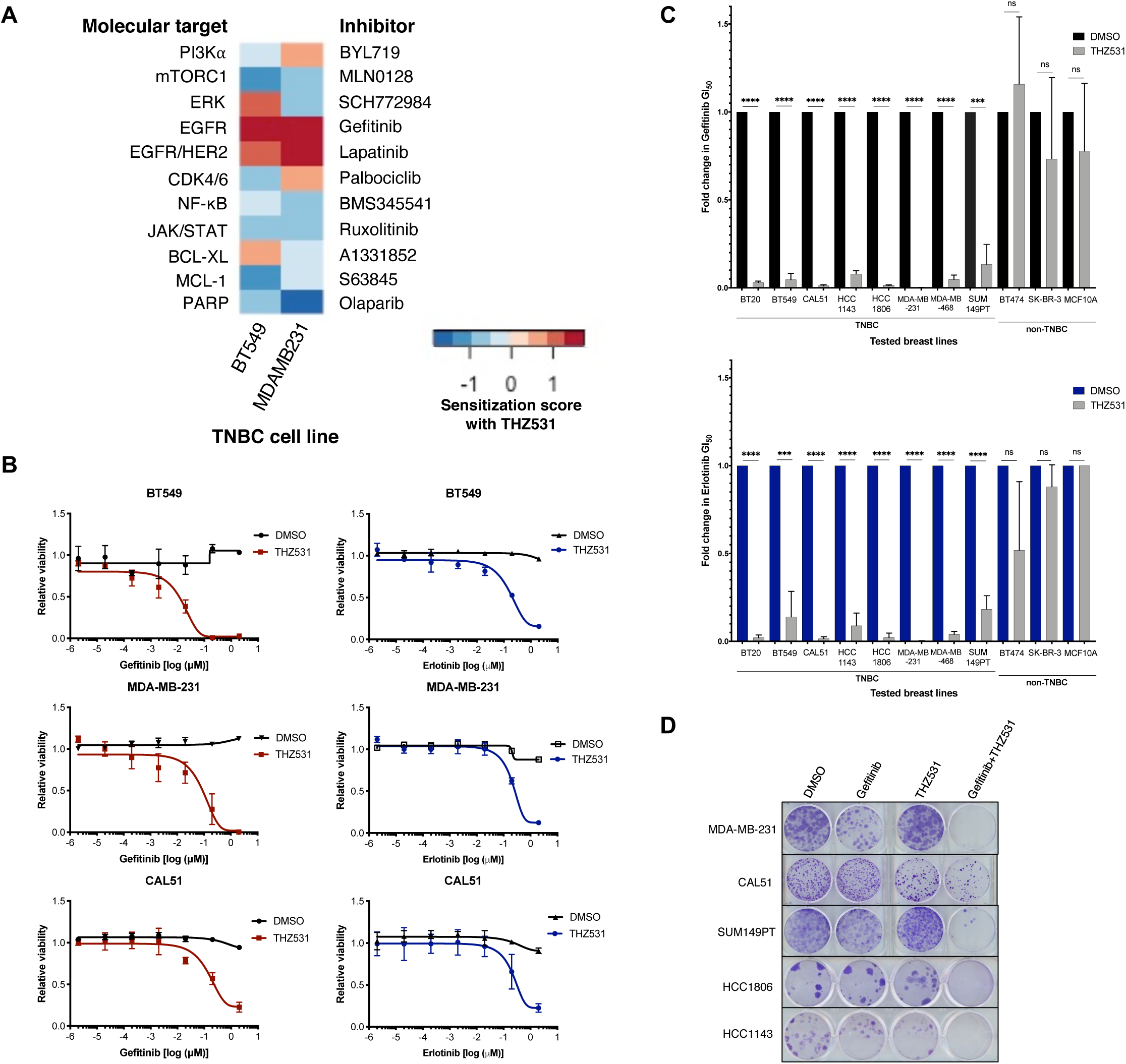
Sensitization to EGFR inhibition with THZ531 in TNBC cells. **(A)** Heatmap depicting the sensitization scores (ratio of GI_50_ values for the indicated drug in the absence versus presence of THZ531, log_10_ transformed) for a panel of inhibitors targeting potential oncogenic drivers in BT549 and MDA-MB-231 TNBC cell lines. (THZ531 background concentrations were 50 nM for BT549 and 200 nM for MDA-MB-231.) **(B)** EGFR inhibitor (geftinib and erlotinib) dose-response curves in BT549, MDA-MB-231 and CAL51 treated with DMSO control or THZ531 in the background (BT549, 50 nM; MDA-MB-231 and CAL51, 200 nM). **(C)** Fold change in 72h GI_50_ values for EGFR inhibitors (gefitinib or erlotinib) in the presence of DMSO control or THZ531 in the background across a panel of breast cancer cell lines. (MCF10A is an immortalized, non-malignant breast epithelial cell line.) Within each cell line, absolute GI_50_ values are normalized to vehicle treatment. (THZ531 background doses: BT549 and SK-BR-3, 50 nM; HCC1143 and HCC1806, 100 nM; BT474 and MDA-MB-468, 150 nM; BT20, CAL51, MCF10A and MDA-MB-231, 200 nM; SUM149PT, 250 nM). ns = not significant, **p ≤ 0.01, ***p ≤ 0.001, ****p ≤ 0.0001 by Student’s t tests; n = 3. Data are mean ± SD of three biological replicates. **(D)** Clonogenic growth assay of TNBC cells (MDA-MB-231, CAL51, SUM149PT, HCC1806, HCC1143) treated with DMSO, gefitinib, THZ531, or geftinib+THZ531 (geftinib at 1 µM; THZ531 doses as in (C)).

### Concurrent EGFR and CDK12 inhibition decreases the levels of key oncogenic proteins in TNBC cells

The global involvement of CDK12 in transcription elongation and multiple post-transcriptional events, including mRNA splicing and intronic polyadenylation (Dubbury et al., 2018; Bartkowiak et al, 2019; Greenleaf, 2019; Choi et al., 2020), prompted us to examine the genes whose expression was selectively affected by the drug combination. Using an unbiased transcriptomic approach, we performed RNA-sequencing in two TNBC cell lines treated with vehicle control, single agents gefitinib and THZ531, and the combination. Consistent with CDK12’s role in transcriptional regulation, THZ531 treatment resulted in the differential expression of thousands of genes. However, relatively few genes were further differentially expressed when gefitinib was added to THZ531 (Figure 2A). This suggested that the synergistic activity of the combined treatment may not be chiefly driven by gene expression changes. In parallel, we evaluated the levels of key oncogenic proteins in TNBC cells treated with vehicle control, single agents, and the combination. In three TNBC cell lines, combined treatment with gefitinib and THZ531 suppressed the levels of MYC, MCL-1, and p300 proteins (Figure 2B). These proteins are notable, as extensive studies have documented their roles as driver oncoproteins in TNBC (DepMap, 2019; Ghandi et al., 2019; Goga et al., 2007; Horiuchi et al., 2012; Hsu et al., 2015; Camarda et al., 2016; Horiuchi et al., 2016; Lee et al., 2017; Klauber-DeMore et al., 2018; Zhao et al., 2018; Rohrberg et al., 2020; Bowling et al., 2021; Goodwin et al., 2015; Merino et al., 2017; Yang et al., 2013).

**Figure 2.**
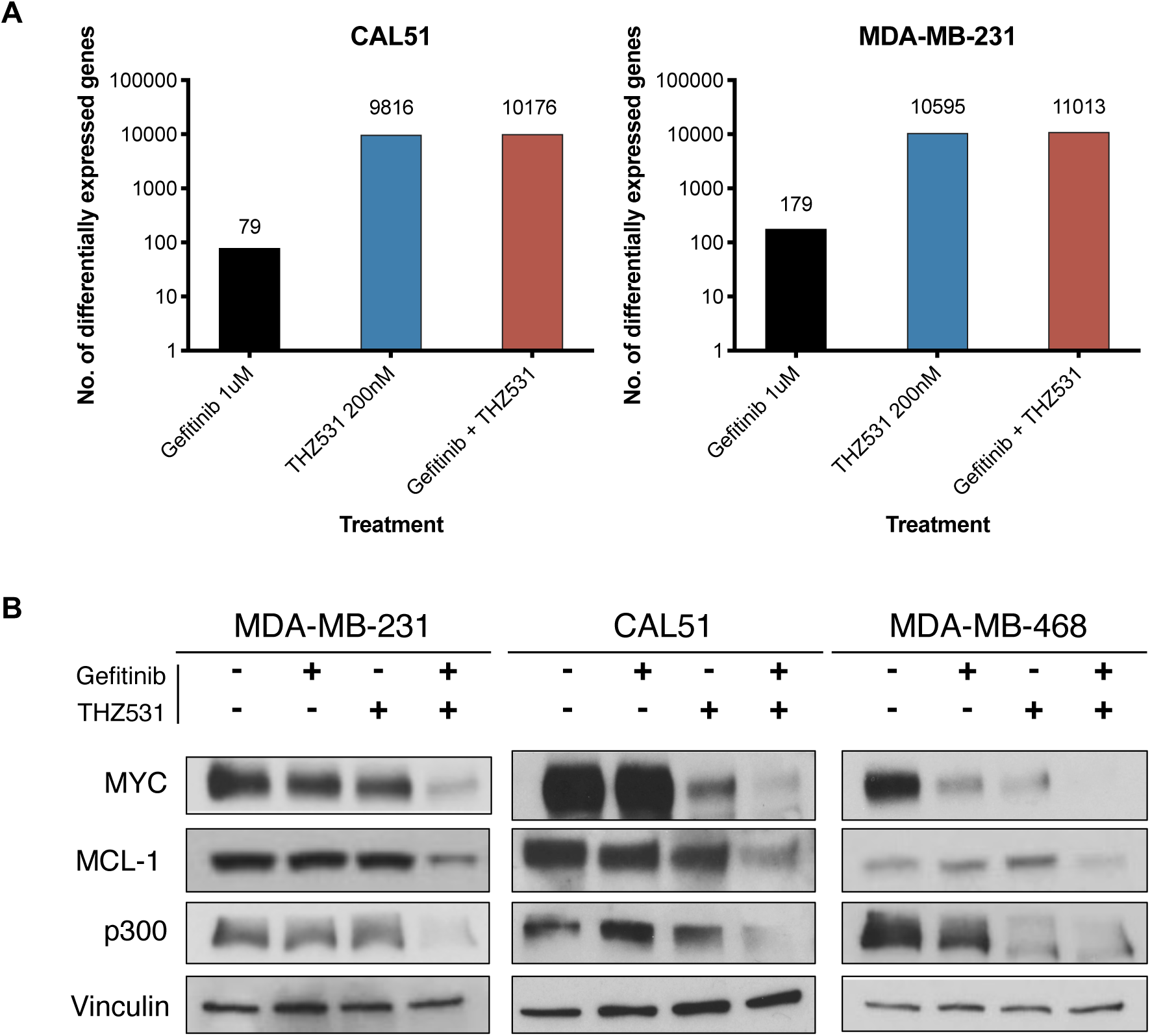
Concurrent EGFR and CDK12 inhibition results in the loss of key oncogenic proteins in TNBC cells. **(A)** Number of differentially expressed genes, assessed by RNA-seq of cells treated with gefitinib (1 µM), THZ531 (200 nM), or the combination, compared to DMSO control, in CAL51 and MDA-MB-231 cells. **(B)** Immunoblot analysis of MYC, MCL-1, p300, and vinculin levels in TNBC cells (MDA-MB-231, CAL51, MDA-MB-468) treated as indicated (geftinib at 1 µM; THZ531 doses as in Figure 1C).

### MYC protein levels are suppressed through both decreased protein translation and increased ubiquitin-proteasome-dependent protein degradation

Given the particularly well-established role of MYC as a driver oncoprotein in TNBC, coupled with the lack of translational therapeutic strategies to inhibit its function in patients, we sought to understand the basis for its loss following co-inhibition of EGFR and CDK12. Specifically, we surveyed each step of MYC biogenesis – transcription, translation, and degradation. Direct analysis of MYC mRNA transcription using quantitative real-time PCR showed increased mRNA expression in gefitinib plus THZ531 treated cells (Figure S3A), a surprising result given the loss of MYC protein with the same combination. Although previous studies suggested that MYC mRNA levels may be acutely suppressed with CDK12 inhibition (Zhang et al, 2016; Krajewska et al., 2019), our data reveal that on the timescale of therapeutic effects, MYC transcript levels are increased and thus cannot account for the observed MYC protein loss.

We next examined the effects of gefitinib and THZ531 on mRNA translation using sucrose density gradient polysome profiling. Polysome profiles showed global reductions in heavy polysomes and enhanced levels of 80S monosomes, consistent with global reduction in mRNA translation in cells treated with THZ531 alone and the drug combination (Figure 3A, boxed polysome fractions). Specifically, the gradient distribution of MYC mRNA also showed reduced association with heavy polysomes, suggesting decreased MYC mRNA translation following treatment with the combination but not the individual drugs (Figure 3A, right panel). We confirmed the decrease in MYC translation using [^35^S]methionine labeling followed by MYC immunoprecipitation in cells treated with gefitinib+THZ531. As observed in the phosphorimage and normalized quantification, [^35^S]methionine incorporation into nascent MYC proteins was considerably decreased in the combination-treated versus the vehicle-treated samples (Figure 3B and S3B), providing direct evidence of MYC translational suppression in the presence of combined gefitinib and THZ531.

**Figure 3.**
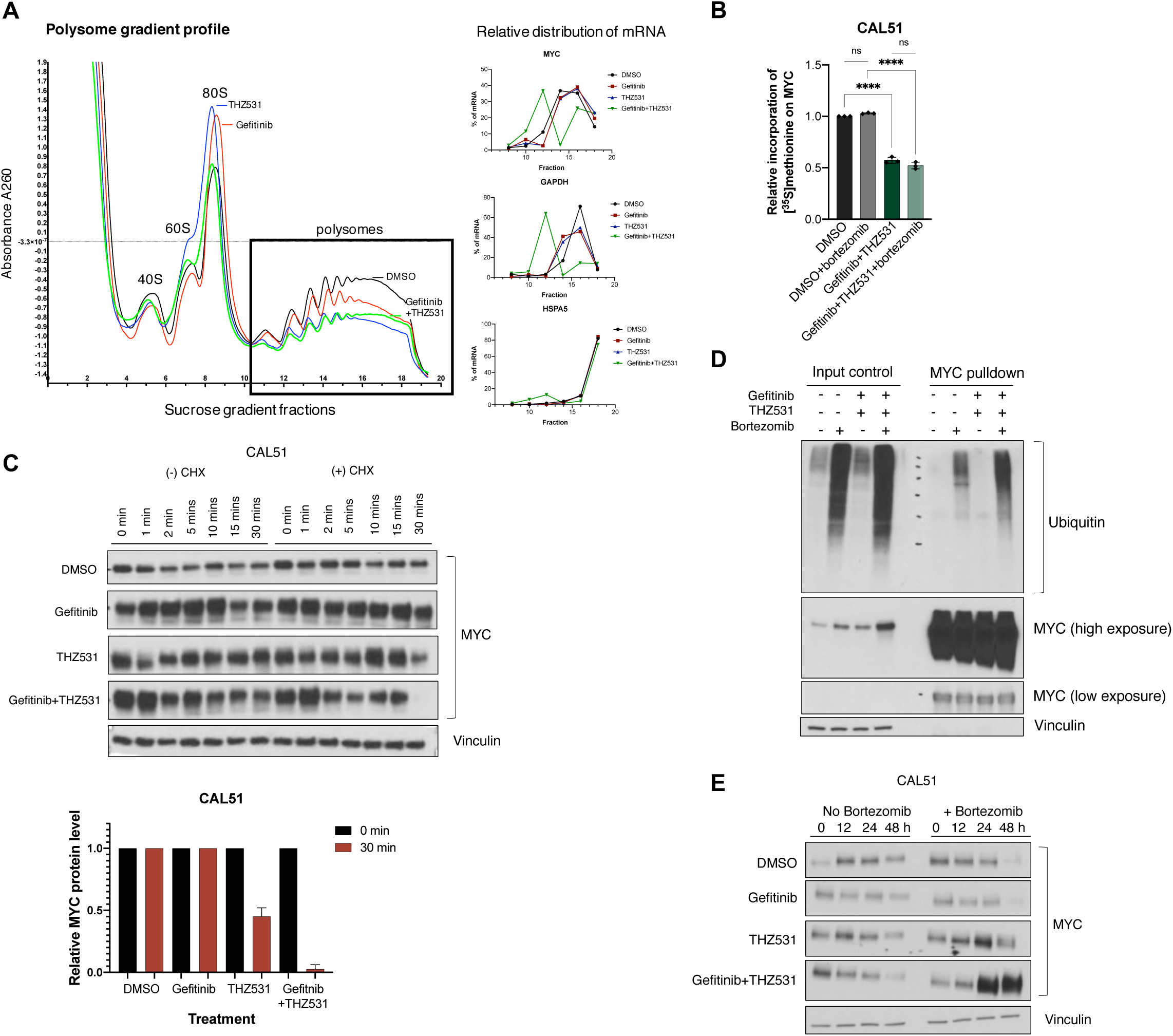
MYC protein loss is driven by decreased protein translation and increased ubiquitin-proteasome-dependent protein degradation. **(A)** Polysome gradient profile in CAL51 cells treated with DMSO, gefitinib (1 µM), THZ531 (200 nM), or gefitinib + THZ531 for 12h. Polysomal fractions highlighted in black box. At right, the relative distribution of MYC, GAPDH, and HSPA5 mRNA levels across the sucrose gradient of the indicated fractions as determined by qRT-PCR. Representative analysis of the polysome distribution of n = 2 independent experiments yielding similar results. **(B)** Relative incorporation of [^35^S]methionine on MYC protein, derived from densitometry quantification of phosphorimage and immunoblot from [^35^S]methionine radioactivity labeling and immunoprecipitation of nascent MYC protein in the absence or presence of proteasome inhibitor, bortezomib (20 nM) in CAL51 cells treated with DMSO or gefitinib (1 µM) + THZ531 (200 nM) for 12h (Fig. S3B). ns = not significant, **p ≤ 0.01, ***p ≤ 0.001, ****p ≤ 0.0001 by Student’s t tests; n = 3. Data are mean ± SD of three biological replicates. **(C)** Immunoblot analysis of MYC and vinculin over a time course as indicated in the absence or presence of cycloheximide (20 µg/mL) in CAL51 cells treated with DMSO, gefitinib (1 µM), THZ531 (200 nM), or gefitinib + THZ531. Relative MYC protein level at time 0 and 30 min with indicated treatment conditions, derived from densitometry quantification of immunoblots from cycloheximide chase experiment of CAL51 cells. Data are mean ± SD of n = 2 independent experiments. **(D)** Immunoblot analysis of ubiquitin, MYC and vinculin on immunoprecipitated MYC proteins and input control in the absence or presence of proteasome inhibitor, bortezomib (20 nM) in CAL51 cells treated with DMSO or gefitinib (1 µM) + THZ531 (200 nM) for 18h. **(E)** Immunoblot analysis of MYC and vinculin over a time course as indicated in the absence or presence of bortezomib (20 nM) in CAL51 cells treated with DMSO, gefitinib (1 µM), THZ531 (200 nM), or gefitinib + THZ531.

To examine MYC stability in the combination treatment protocol, we performed a cycloheximide chase experiment. MYC protein stability was unaffected by gefitinib, but slightly decreased with THZ531 alone and to a greater extent with the combination treatment (Figure 3C and S3B). We further investigated the canonical MYC protein degradation pathway by the ubiquitin-proteasome system. Following MYC protein immunoprecipitation, we observed a distinctive increase in MYC ubiquitination in cells treated with gefitinib plus THZ531 (Figure 3D and S3D). Consistent with these data, proteasome inhibition by bortezomib rescued the decline in MYC protein affected by the drug combination, indicating proteasome-dependent degradation of MYC (Figure 3E). Together, these data demonstrate that the cumulative loss of MYC protein following combined EGFR and CDK12 inhibition resulted from suppressed MYC translation and increased MYC protein degradation.

### EGFR and CDK12 inhibition synergistically decrease MYC protein stability through their regulation of 4E-BP1 phosphorylation

It was recently reported that CDK12 acts as a positive regulator of cap-dependent translation through direct phosphorylation of the mRNA 5’ cap-binding repressor, 4E-BP1 (Choi et al., 2019). This led us to consider whether the suppression of MYC protein translation following combined EGFR and CDK12 inhibition occurs via the dephosphorylation of 4E-BP1. In line with this hypothesis, we observed significant dephosphorylation of the four well-established 4E-BP1 phosphosites (T37, T46, S65, and T70) with the drug combination (Figure 4A). This indicates that combined CDK12 and EGFR inhibition prevented phosphorylation of 4E-BP1, thus retaining cap-binding on mRNA and suppressing cap-dependent mRNA translation.

**Figure 4.**
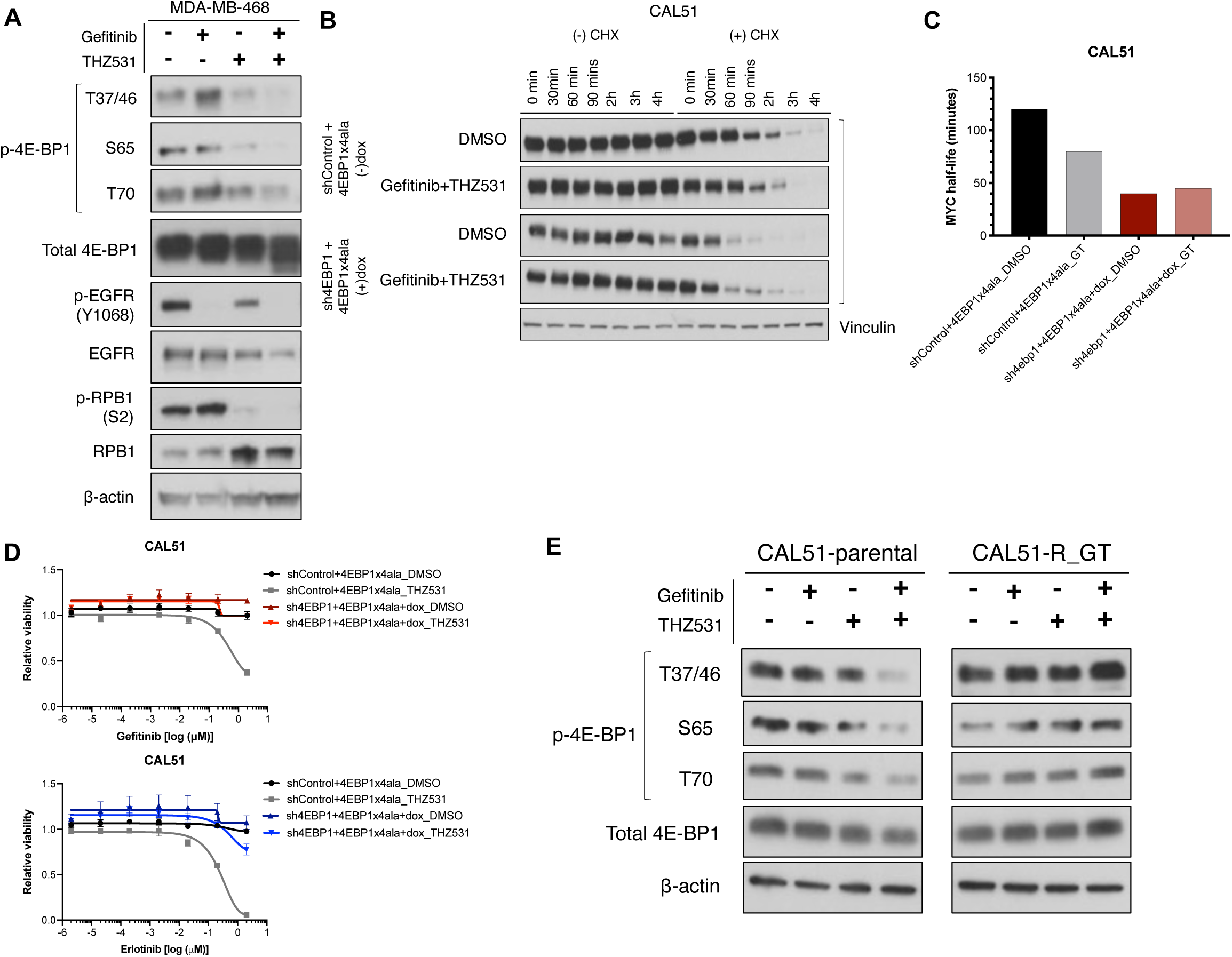
Combined EGFR and CDK12 inhibition destabilizes MYC protein and drives cell death through regulation of 4E-BP1 phosphorylation. **(A)** Immunoblot analysis of 4E-BP1 phosphorylation at T37/46, S65, and T70, total 4E-BP1, phospho-EGFR (Y1068), total EGFR, phospho-RPB1 (S2), total RPB1, and β-actin in MDA-MB-468 cells treated with DMSO, gefitinib (1 µM), THZ531 (150 nM), or gefitinib + THZ531 for 24h. **(B)** Immunoblot analysis of MYC and vinculin over a time course as indicated in the absence or presence of cycloheximide (20 µg/mL) in CAL51 cells expressing a doxycycline-inducible, non-phosphorylatable 4E-BP1 mutant and a 3’ UTR-targeted sh4EBP1 construct (to remove endogenous 4E-BP1), treated with DMSO or gefitinib (1 µM) + THZ531 (200 nM). **(C)** MYC protein half-life derived from densitometry quantification of immunoblots from cycloheximide chase experiment of CAL51 4E-BP1-manipulated derivatives in (B). **(D)** EGFR inhibitor (geftinib and erlotinib) dose-response curves in CAL51 4E-BP1-manipulated derivatives treated with or without THZ531 (200 nM) in the background. **(E)** Immunoblot analysis of 4E-BP1 phosphorylation at T37/46, S65 and T70, total 4E-BP1, and β-actin in gefitinib + THZ531 (GT)-resistant and parental CAL51 cells treated with DMSO, gefitinib (1 µM), THZ531 (200 nM), or gefitinib + THZ531 for 12h.

Work in model systems has demonstrated that defects in translation can lead to the suppression of protein stability (Stein and Frydman 2019; Sontag et al., 2017; Brandman and Hedge, 2016; Joazeiro, 2019). To understand whether 4E-BP1-dependent suppression of cap-dependent translation drives the reduction in MYC stability, we used a 3’ UTR-targeted shRNA to suppress the expression of endogenous 4E-BP1, then simultaneously expressed a dominant negative, non-phosphorylatable mutant of 4E-BP1 (T37A, T46A, S65A, T70A) under doxycycline-responsive control (Figure S4A). In this configuration, MYC half-life was substantially suppressed, indicating that suppression of MYC translation is alone sufficient to destabilize the protein. Further, we observed that the non-phosphorylatable mutant 4E-BP1 blocked the ability of gefitinib + THZ531 to further destabilize MYC, suggesting that the drug combination destabilizes MYC through its effects on 4E-BP1 phosphorylation (Figure 4B and 4C).

Beyond its effects on MYC, the overall cytotoxic effect of the combination therapy was also dependent on 4E-BP1, as replacement of endogenous 4E-BP1 with the non-phosphorylatable mutant blocked the cellular response to the drug combination (Figure 4D). Consistent with this finding, TNBC cells cultured chronically in media containing gefitinib and THZ531 until they developed resistance were insensitive to drug-induced suppression of 4E-BP1 phosphorylation (Figure 4E, S4B and S4C). Together, these findings demonstrate that combined EGFR and CDK12 inhibition leads to MYC destabilization and cell death through the cooperative regulation of 4E-BP1 activity.

### CNOT1 is required for combination therapy-induced MYC translational suppression, MYC degradation, and cell death

To gain further insight into the mechanisms underlying the biological activity of the combination therapy, we performed genome-wide CRISPR/Cas9-based loss-of-function screens in cells treated with vehicle control or gefitinib + THZ531 (Figure 5A). We focused our analysis on genes whose knockouts were enriched in the combination-treated arm, as genes scoring in this group can be interpreted as being required for the full activity of the combination therapy. Analysis of the screen revealed that multiple CNOT family gene knockouts were enriched in cells treated with gefitinib + THZ531 (Figure 5B). The CCR4-NOT complex, which is comprised of multiple CNOT family proteins, is reported to be functionally involved in post-transcriptional mRNA deadenylation, translational quality control, and protein ubiquitylation (Shirai et al., 2014; Miller and Reese, 2012). Among the CNOT family genes, CNOT1, a central scaffolding component of the CCR4-NOT complex, was the top scoring gene in our screen. CNOT1 knockout led to a nearly complete rescue of the cooperativity between EGFR and CDK12 inhibitors as well as the overall toxicity of the drug combination (Figure 5C and 5D). Molecularly, CNOT1 loss rendered cells insensitive to drug-induced loss of both 4E-BP1 phosphorylation and consequent MYC and MCL-1 protein loss (Figure 5E). Further, CNOT1 protein expression was lost in cells with naturally evolved resistance to the combination therapy (Figure 5F), suggesting that it may be responsible for the resistance of these cells to drug-induced 4E-BP1 dephosphorylation and death. Together, the results of this study are summarized in Figure 5G (Mathys et al., 2014; Chen et al., 2014; Cooke et al., 2010; Kamenska et al, 2016; Ozgur et al; 2015; Miller and Reese, 2012; Panasenko and Collart, 2011).

**Figure 5.**
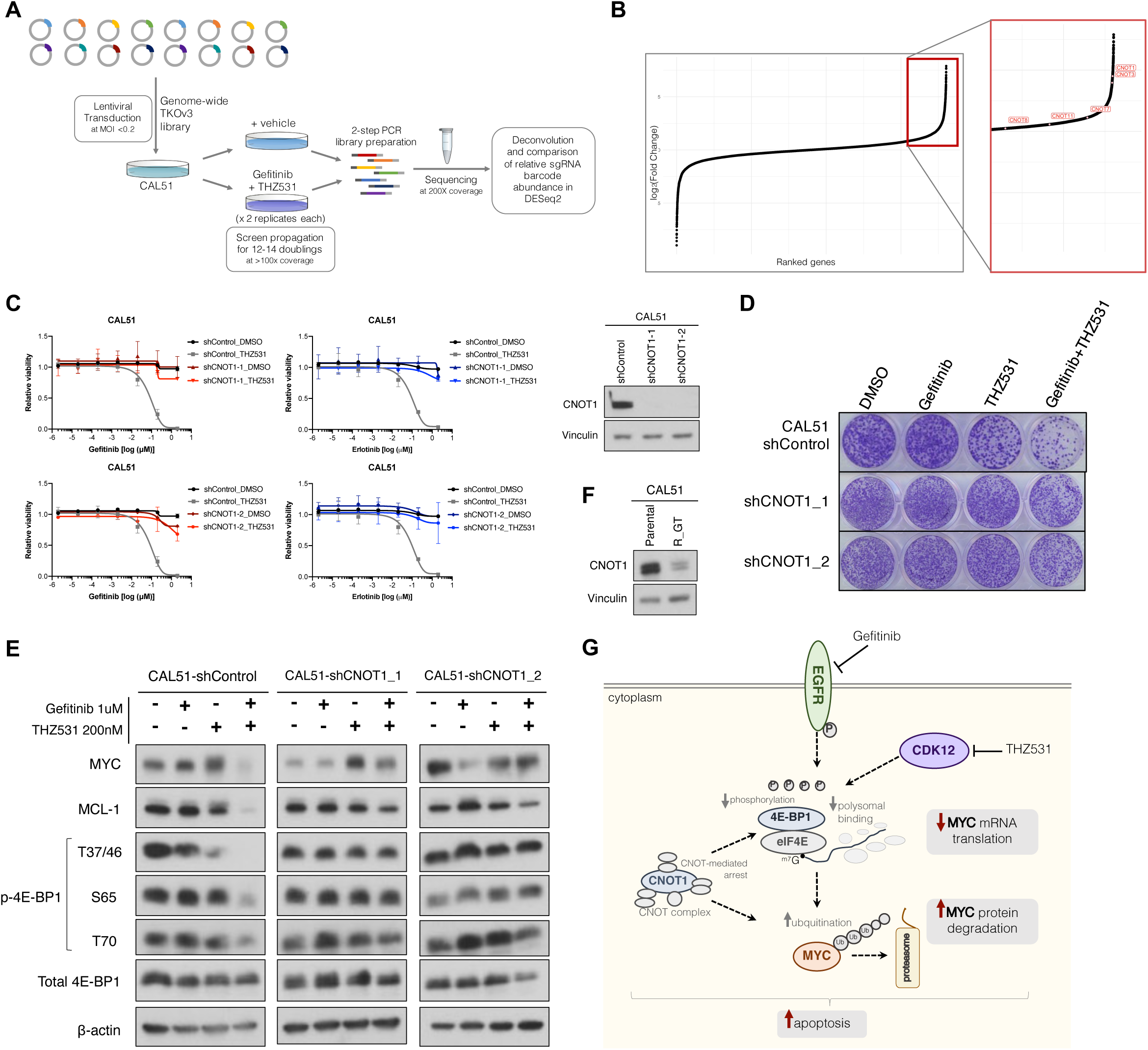
CNOT1 is required for combination therapy-induced translational suppression, MYC degradation, and cell death. **(A)** Schematic overview of genome-wide CRISPR positive selection screen. **(B)** Gene-level representation of screen results, ranked by log_2_(fold change). Cluster of CNOT family genes (highlighted in red) are enriched among those knockouts (boxed) enriched in the gefitinib+THZ531 combination-treated population. **(C)** EGFR inhibitor (geftinib and erlotinib) dose-response curves in CAL51 cells expressing shControl or each of two independent shRNAs targeting CNOT1, treated with or without THZ531 200 nM in the background. Immunoblot analysis of CNOT1 and vinculin in CAL51 cells expressing the indicated shRNAs (right). **(D)** Clonogenic growth assay of CAL51 cells expressing the indicated shRNAs and treated with DMSO, gefitinib (1 µM), THZ531 (200 nM), or geftinib + THZ531. **(E)** Immunoblot analysis of MYC, MCL-1, phospho-4E-BP1 (T37/46, S65, T70), total 4E-BP1 and β-actin in CAL51 expressing the indicated shRNAs and treated with DMSO, gefitinib (1 µM), THZ531 (200 nM), or geftinib + THZ531 for 24h. **(F)** Immunoblot analysis of CNOT1 and vinculin in GT-resistant and DMSO-control CAL51 cells. **(G)** Summarized mechanistic model of cooperative regulation of MYC biogenesis with EGFR and CDK12 inhibition in TNBC via 4E-BP1 hypo-phosphorylation and CNOT-complex mediated processes in TNBC cells.

## DISCUSSION

The discovery, over a decade ago, of EGFR hyperactivation in TNBC and its correlation with aggressive disease, chemoresistance, and poor prognosis positioned EGFR as a prime therapeutic target in this disease subset. However, the subsequent observation of poor clinical responses to EGFR inhibitors in TNBC patients dampened enthusiasm for this therapeutic target and raised a fundamental question: Is EGFR a driver of TNBC pathogenesis whose importance is obscured by mechanisms of intrinsic resistance, or is it simply a bystander signaling event (Ali and Wendt, 2017)? Here, we resolve this question, demonstrating that the inhibition of CDK12 reveals an exquisite dependence of diverse TNBC models on EGFR signaling. Further, these studies reveal that EGFR and CDK12 cooperate to drive TNBC not through transcriptional regulation, but rather by promoting the translation and associated stabilization of key driver oncoproteins, including MYC (Goga et al., 2007; Horiuchi et al., 2012; Hsu et al., 2015; Camarda et al., 2016; Horiuchi et al., 2016; Klauber-DeMore et al., 2018; Zhao et al., 2018; Rohrberg et al., 2020; Bowling et al., 2021). As such, the coupled nature of protein synthesis and stability, which has been well-explored in model systems (Stein and Frydman 2019; Sontag et al., 2017; Brandman and Hedge, 2016; Joazeiro, 2019), is shown here, for what we believe to be the first time, to underlie the therapeutic activity of a promising anti-cancer strategy.

Several key open questions remain. First, while the drug combination under study clearly functions through the modulation of 4E-BP1-regulated, coupled protein translation and stability, the relative contributions of decreased oncoprotein synthesis versus decreased stability to the observed toxicity have not been clarified. Further, the extent to which EGFR- and/or CDK12-regulated transcriptional events or proteasome modulation may template the observed mechanism of action has not been resolved. Second, a number of reports have described cooperativity between EGFR blockade and inhibition of receptor tyrosine kinase (RTK)-PI3K-mTOR signaling in TNBC (El Guerrab et al., 2020; You et al., 2018; Simiczyjew et al., 2018; Verma et al., 2017; Tao et al., 2014; Sohn et al., 2014). A recent report also demonstrated that the effects of EGFR inhibition can be potentiated through blockade of Elongator complex-mediated MCL-1 translation (Cruz-Gordillo et al., 2020). While these studies revealed sensitizing effects that appear to be significantly less profound, and more heterogeneous, than those reported here, it remains to be determined if these processes regulate, or are regulated by, CDK12. Third, although this study identifies MYC, MCL-1, and p300 protein loss as likely key events downstream of combined EGFR and CDK12 inhibition, unbiased proteomic approaches may reveal additional TNBC driver oncoproteins that are similarly affected. Fourth, the precise mechanisms by which the CCR4-NOT impacts EGFR- and CDK12-regulated 4E-BP1 activity and downstream oncoprotein stability remain to be defined. The CCR4-NOT complex has been shown to regulate translation by interacting with translational repressors such as eIF4E and DDX6 and blocking decapping machinery (Mathys et al., 2014; Chen et al., 2014; Cooke et al., 2010; Kamenska et al, 2016; Ozgur et al; 2015). Further, it also functions in the ubiquitination of nascent, translationally arrested polypeptides and the maintenance of 26S proteasome integrity (Miller and Reese, 2012; Panasenko and Collart, 2011), suggesting that its regulatory roles in the phenomena under study here may be multifactorial. Finally, the full therapeutic potential of combined EGFR and CDK12 inhibition has not been evaluated in preclinical animal models because THZ531 is not amenable to *in vivo* administration.

In sum, by revealing a long debated EGFR dependence in TNBC, we have identified a therapeutic approach that functions through a novel, unexpected mechanism of action and holds promising translational potential for the treatment of this difficult-to-treat disease subtype.

## ACKNOWLEDGEMENTS

We thank the members of the K.C.W. and C.V.N. laboratories as well as Sarah Sammons and James Alvarez for their valuable support and feedback on the manuscript. We thank Arno Greenleaf for helpful discussions and for providing CDK12^AS^-expressing cells. We thank Tim Haystead and David Loiselle for facilitating the [^35^S]methionine labeling experiment. This work was supported by Duke University School of Medicine start-up funds and support from the Duke Cancer Institute (K.C.W.), an NIH award (R01CA207083 to K.C.W.), DoD Breast Cancer Research Program awards (W81XWH1910414 and W81XWH1610703 to K.C.W.), and the Agency for Science, Technology and Research, Singapore (NSS-PhD to H.X.A).

## AUTHOR CONTRIBUTIONS

Conceptualization, H.X.A., C.V.N., K.C.W.; Methodology, H.X.A., T.E.R., C.V.N., and K.C.W., Investigations, H.X.A., N.S., X.L.D., L.C.B., Q.C.; Formal Analysis; H.X.A., I.C.M., A.B., J.L., J.S. and J.P.H.; Writing – Original Draft, H.X.A. and K.C.W.; Writing – Review & Editing, all authors; Visualization, H.X.A.; Supervision, T.E.R., C.V.N., and K.C.W.; Funding Acquisition, H.X.A., C.V.N., and K.C.W.

## STAR METHODS

Detailed methods are provided in the online version of this paper and include the following:

- Key resources table
- Contact for reagents and resource sharing
- Experimental model
- Method details
- Quantification and statistical analysis
- Data and software availability

## SUPPLEMENTAL INFORMATION

Supplemental Information includes three figures and four tables and can be found with the online version of this paper.

## SUPPLEMENTAL INFORMATION

### Supplemental Tables

**Table S1A.** CAL51 RNA-seq counts with ERCC spike-in normalization

**Table S1B.** MDA-MB-231 RNA-seq counts with ERCC spike-in normalization

**Table S2.** Raw counts for TKOv3 positive selection screen

**Table S3.** Analyzed screen gene list for DESeq2 comparison analysis GT vs DMSO

### Supplemental Figures

**Figure S1.**
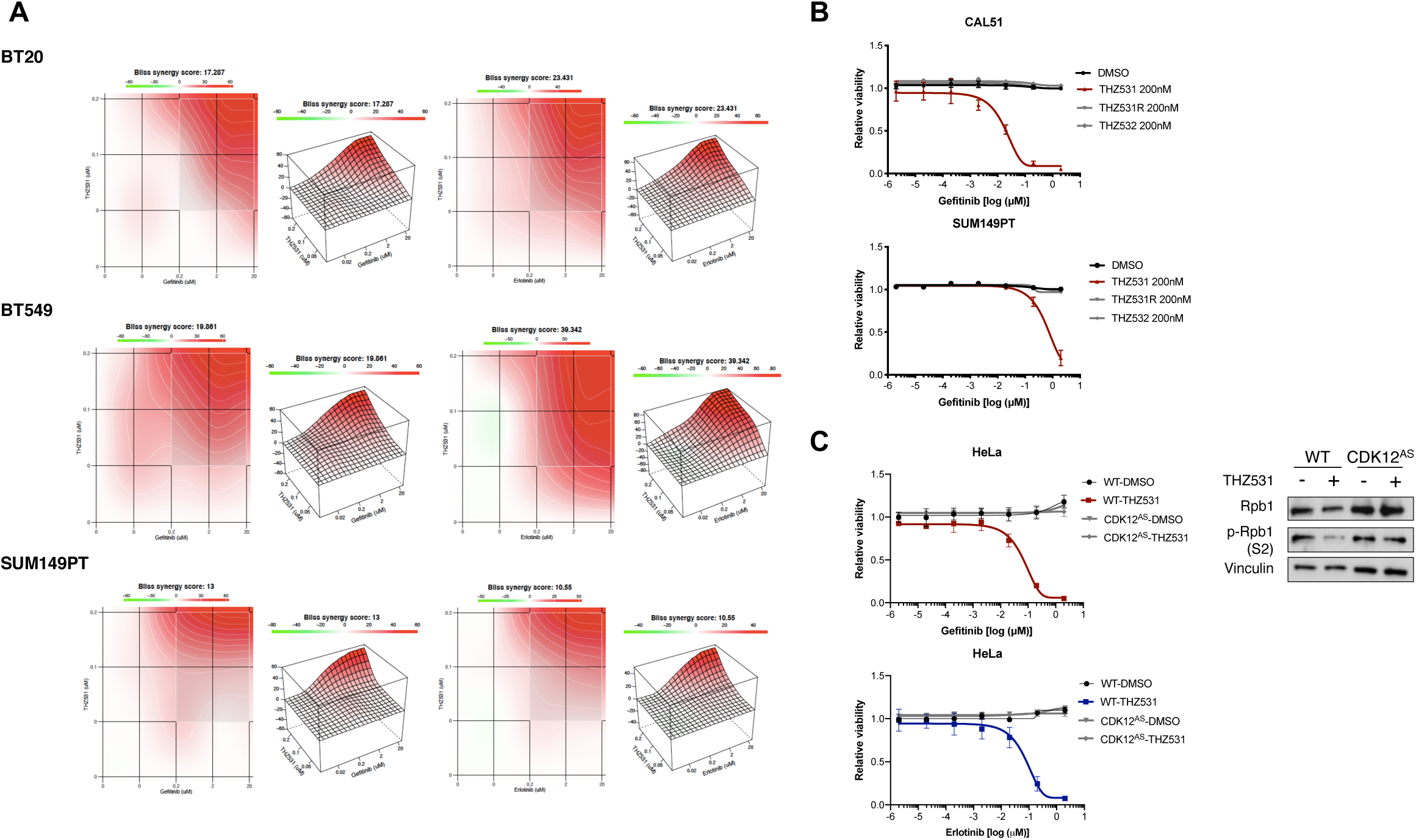
Associated with Figure 1. **(A)** Bliss synergy scores confirmed synergistic activity (score > 10) between EGFR inhibitors (gefitinib and erlotinib) and THZ531 in three representative TNBC cell lines (BT20, BT549, and SUM19PT). **(B)** EGFR inhibitor (gefitinib and erlotinib) dose-response curves in two representative TNBC cell lines (CAL51 and SUM149PT) treated with DMSO, THZ531, or either a stereoisomer (THZ531R) or a derivative of THZ531 (THZ532) that lack the ability to inhibit CDK12/13 in the background. **(C)** EGFR inhibitors (gefitinib and erlotinib) dose-response curves in HeLa cells expressing the CDK12^AS^ mutant (F813G), which is not inhibited by THZ531, relative to wildtype (WT) cells. Immunoblot analysis of phospho-RPB1 (Ser2), total RPB1, and vinculin in WT and CDK12^AS^ HeLa cells.

**Figure S3.**
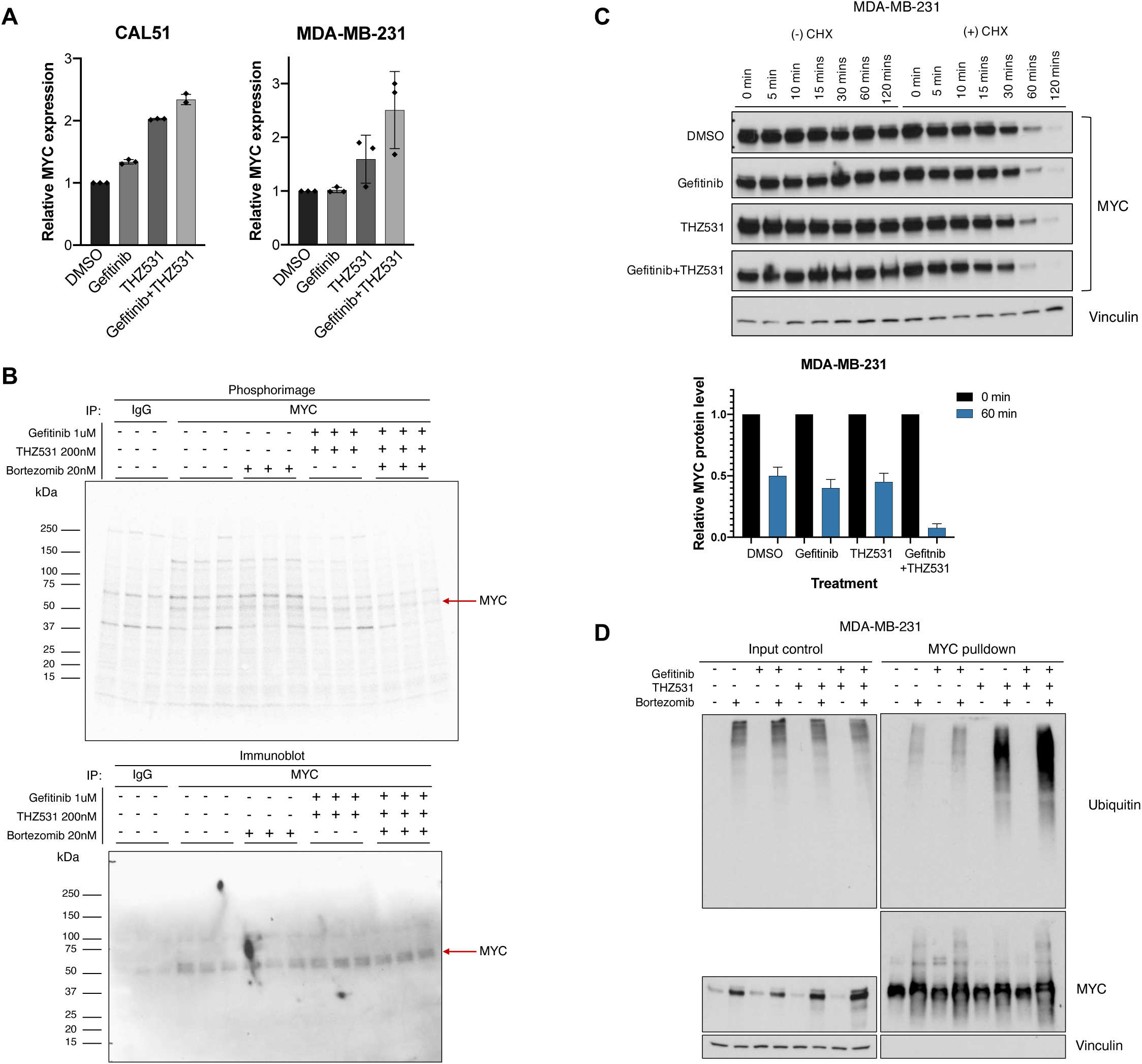
Associated with Figure 3. **(A)** Analysis of relative MYC mRNA expression by qRT-PCR in two representative TNBC cell lines (CAL51 and MDA-MB-231) treated with DMSO, gefitinib (1 μM), THZ531 (200 nM), or geftinib (1 μM + THZ531 200 nM) for 12h. **(B)** Phosphorimage (top) of [^35^S]methionine incorporation radioactivity analysis and immunoblot analysis of MYC (bottom) with immunoprecipitation of nascent MYC protein and IgG isotype control in the absence or presence of proteasome inhibitor, bortezomib (20 nM) in CAL51 cells treated with DMSO or gefitinib (1 µM) + THZ531 (200 nM) for 12h. **(C)** Immunoblot analysis of MYC and vinculin over a time course as indicated in the absence or presence of cycloheximide (20 μg/mL) in MDA-MB-231 cells treated with DMSO, gefitinib (1μM), THZ531 (200 nM), or gefitinib + THZ531. Relative MYC protein level at time 0 and 60 min with indicated treatment conditions, derived from densitometry quantification of immunoblots from cycloheximide chase experiment of MDA-MB-231 cells. Data are mean ± SD of n = 2 independent experiments. **(D)** Immunoblot analysis of ubiquitin, MYC, and vinculin on immunoprecipitated MYC proteins and input control in the absence or presence of the proteasome inhibitor, bortezomib (20 nM) in MDA-MB-231 cells treated with DMSO, gefitinib (1 μM), THZ531 (200 nM), or gefitinib + THZ531 for 18h.

**Figure S4.**
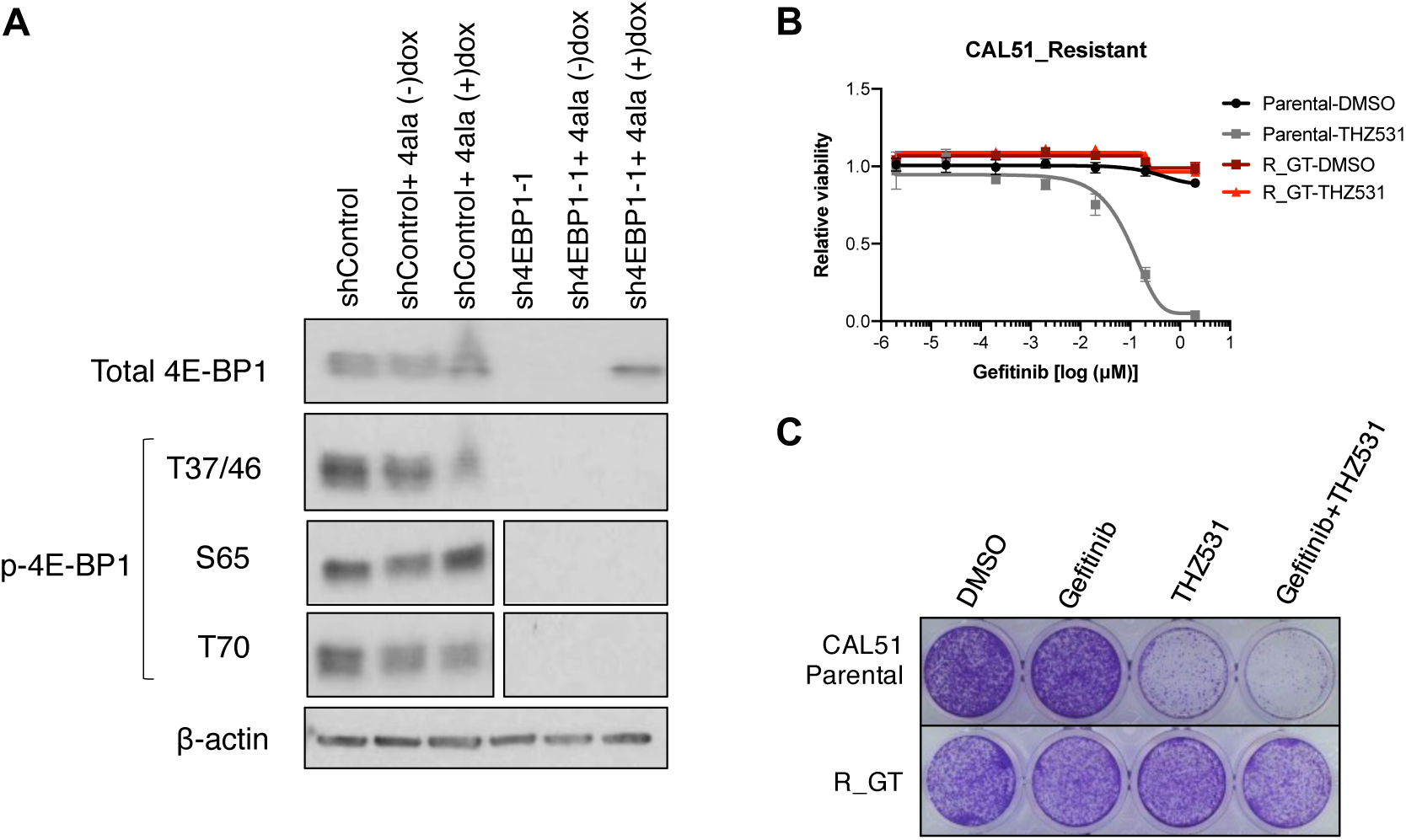
Associated with Figure 4. **(A)** Immunoblot analysis of 4E-BP1 phosphorylation at T37/46, S65 and T70, total 4E-BP1, and β-actin in CAL51 expressing doxycycline-inducible non-phosphorylatable 4E-BP1 mutant and shControl or sh4EBP1. **(B)** Gefitinib dose-response curves in gefitinib + THZ531 (GT)-resistant and parental CAL51 cells. **(C)** Clonogenic growth assay of GT-resistant and parental CAL51 cells treated with DMSO, gefitinib (1 μM), THZ531 (200 nM) or geftinib + THZ531.

## Key Resources Table

### Experimental models

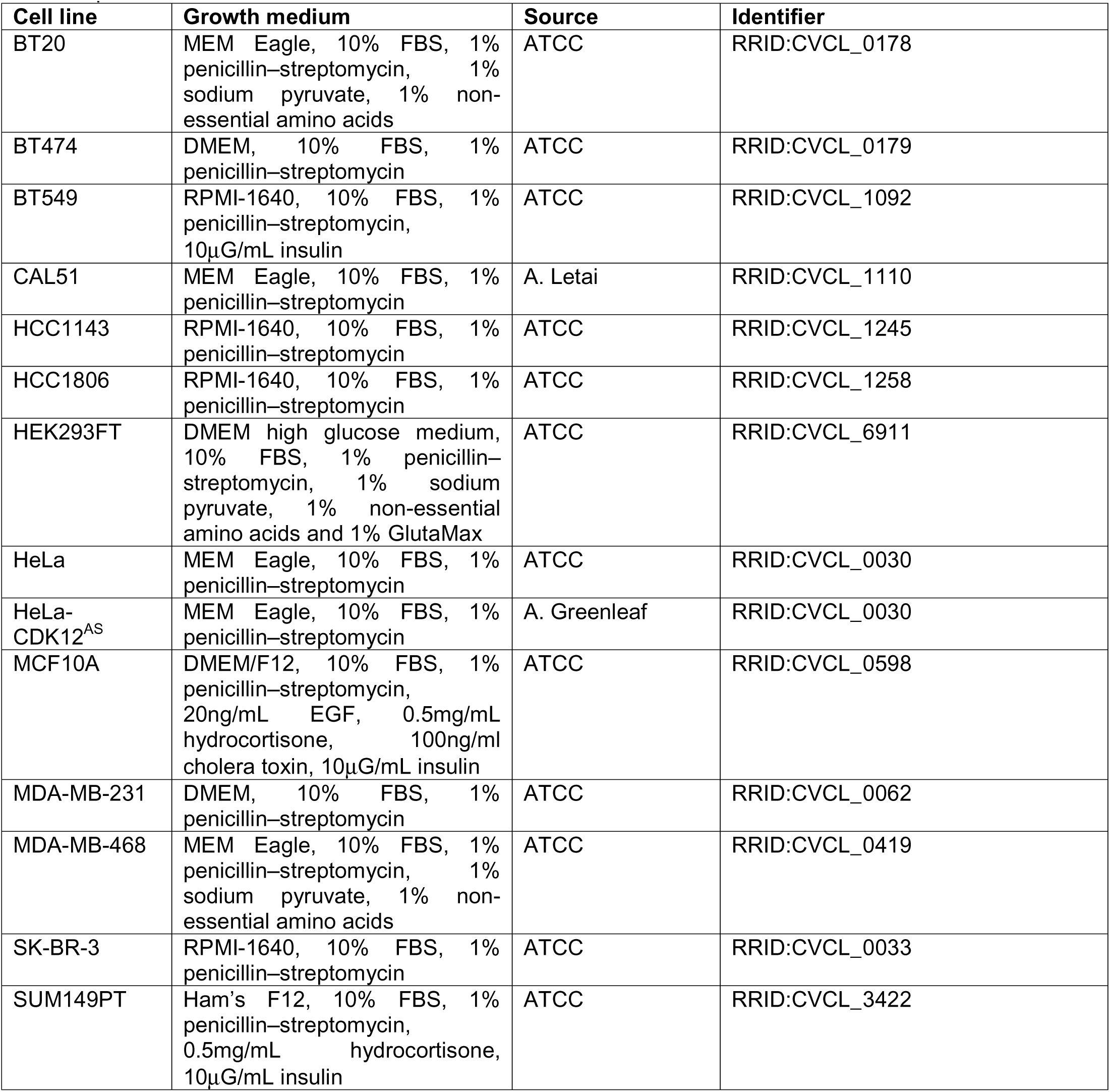
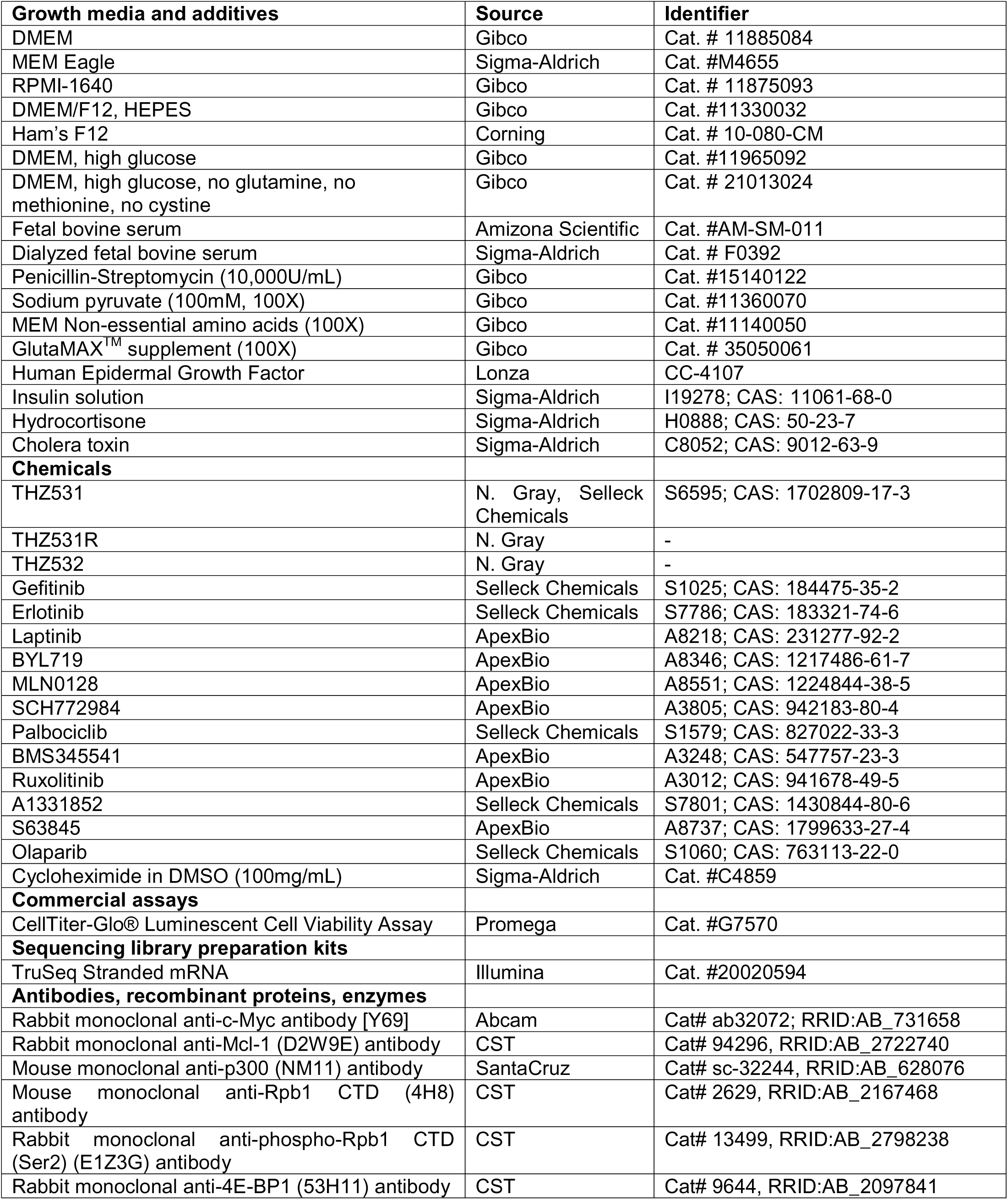
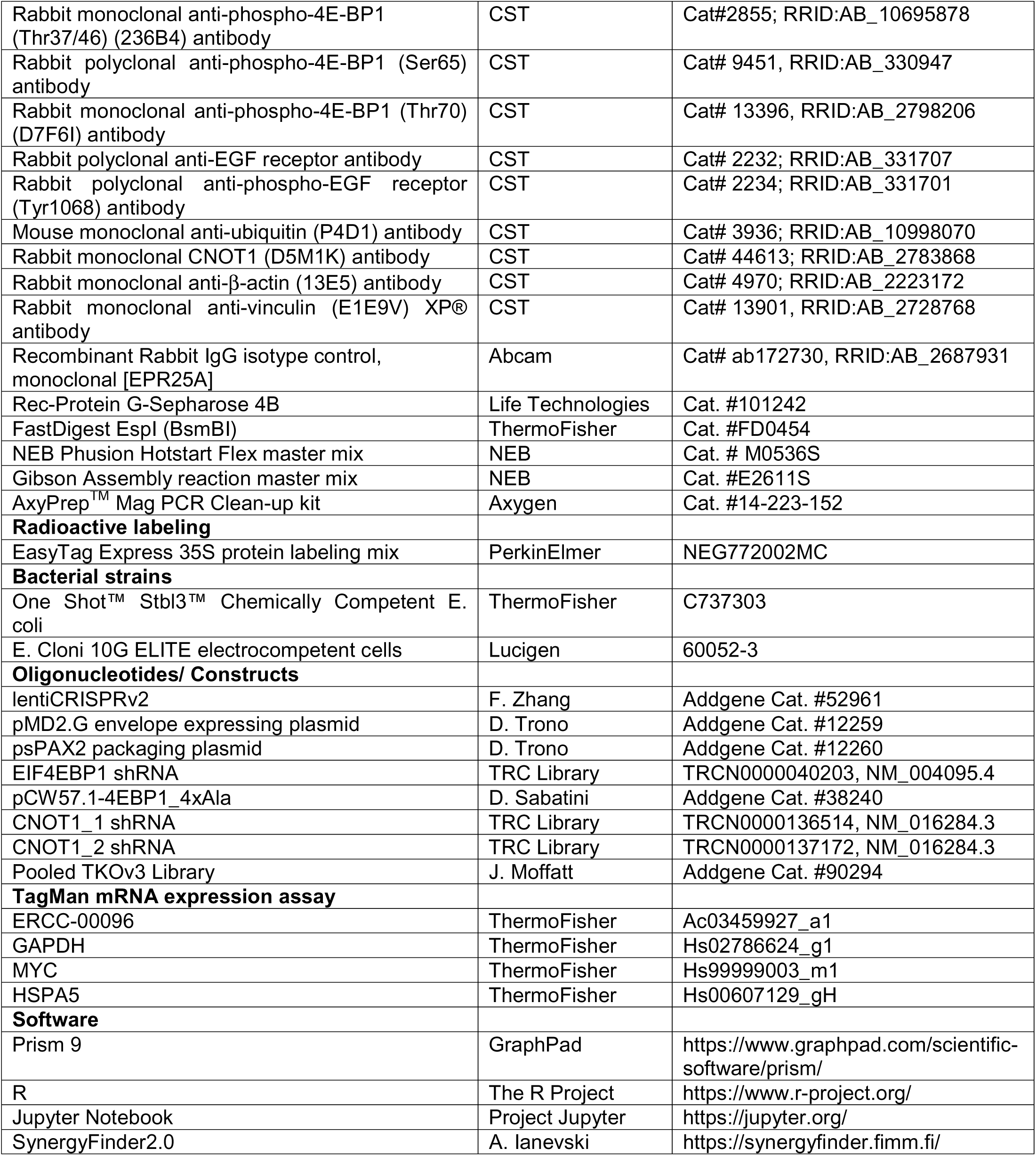

## CONTACT FOR REAGENT AND RESOURCE SHARING

Further information and requests for resources and reagents should be directed to and will be fulfilled by the Lead Contacts, Christopher Nicchitta and Kris Wood.

## EXPERIMENTAL MODEL

### Cell lines, reagents and inhibitors

BT20, BT474, BT549, CAL51, HCC1143, HCC1806, HeLa, MCF10A, MDA-MB-231, MDA-MB-468, SK-BR-3, SUM149PT cell lines were purchased from Duke University Cell Culture Facility (CCF) or American Type Culture Collection (ATCC). CAL51 was gifted by Anthony Letai (Dana Farber Cancer Institute) and HeLa-CDK12^AS^ was kindly gifted by Arnold Greenleaf (Duke University). All cell lines were authenticated using short tandem repeat (STR) profiling by the Duke University DNA Analysis Facility and tested negative for mycoplasma contamination using MycoAlert^TM^ PLUS Mycoplasma Detection kit (Lonza). All cell lines were cultured at 37°C in 5% CO2. See Key Resources Table for specific culture media.

Drugs were purchased from SelleckChem or Apexbio Technology. THZ531R and THZ532 were generously gifted by Nathaniel Gray (Harvard University, Dana-Farber Cancer Institute).

### Drug-resistant CAL51 cell line

To evolve resistance to the gefitinib+THZ531 combination in-vitro, CAL51 cells were exposed to the combined drugs with increasing concentrations, starting with a dose close to 25% of cell viability. As CAL51 cells were insensitive to gefitinib, an arbitrary starting dose of 500nM was selected, while 50 nM of THZ531 was used. The growth rate was monitored by cell counts with passaging every 3-5 days. Once growth rate was stabilized, the concentration of each drug was increased singly each time till maximal preset synergistic dose of 1 μM gefitinib and 200 nM THZ531 to develop CAL51-R_GT. A paired vehicle control was cultured with DMSO-containing media in parallel (CAL51-R_DMSO). Resistant cells were achieved over 8 weeks with gradual dose increments.

## METHOD DETAILS

### GI_50_ and sensitization assay

Cells were seeded in 96-well plates at density of 3000 – 5000 cells per well and treated with a 10-fold serial dilution of indicated drug. Calculated drug dilution series yield final drug concentrations starting with vehicle (DMSO) at 0, 0.000002, 0.00002, 0.0002, 0.002, 0.02, 0.2, 2 μM. CellTiter-Glo luminescent viability assay (Promega) was used to measure cell viability after 72h drug incubation. Luminescence from each specific well of each plate were measured using Tecan plate reader (Infinite M1000 PRO). Each treatment condition was performed in triplicates per plate and represented by three independent experiments. Relative viability was calculated by normalizing raw luminescence values to vehicle-treated wells. GI_50_ values were considered as the dose at which cell viability equates to 50% of DMSO-treated viability and determined by fitting each individual experiment to a 4-parameter logistic drug-response curve using GraphPad/Prism9 software.

For two-drug combinations, the concentration of a second background drug was kept constant across all wells. Sensitization scores were calculated using GI_50_ values with vehicle versus the second background drug as fold change and log_10_ transformed, thus sensitization scores >0 will indicate increased sensitivity to the first serially diluted drug. GI_50_ assays were first performed singly to obtain dose-response curves for THZ531 with each cell line. Background doses for THZ531 were then chosen based on the curves at doses yielding no less than 80% viability to ensure adequate cellular representation of response to first serially diluted drug.

Background doses of THZ531 were: 50 nM for BT549 and SK-BR-3, 100 nM for HCC1143 and HCC1806, 150 nM for BT474 and MDA-MB-468; 200 nM for BT20, CAL51, MCF10A and MDA-MB-231, 250 nM for SUM149PT.

### Bliss synergy score calculation

To quantitatively assess synergy, GI_50_ assays were first performed for each inhibitor (e.g. gefitinib or erlotinib) with a range of four or more fixed concentrations of the second background drug (e.g. THZ531 0, 50, 100, 200 nM). Relative cell viability was calculated as described earlier. Data was tabulated according to SynergyFinder 2.0 User Documentation and uploaded on the web application for analysis (Ianevski et al., 2020). Four-parameter logistic regression (LL4) was selected for curve-fitting algorithm and outlier detection was turned on (Ianevski et al., 2019). Bliss method was selected for synergy calculation with the ‘Correction’ option switched on to eliminate detected outlier and apply a base-line correction method on the single drug dose-responses.

### Clonogenic growth assay

To measure longer term effect of inhibitors and combination on cell growth, cells were seeded at 500 cells/well in 12-well tissue culture plates or 1000 cells/well in 6-well tissue culture plates in complete media. 24h after seeding, media were aspirated, and drugs were added into fresh media to each specific wells. Media and drugs were refreshed every 5 days, and assays were cultured for 10 to 15 days. Drug media were then removed, and plates were fixed and stained with 0.5%w/v crystal violet in 80%v/v methanol solution for 20 min at room temperature. Plates were rinsed with distilled water and scanned.

### Immunoblotting and antibodies

Immunoblotting was performed as previously described (Singleton et al., 2017). Briefly, cells were resuspended in RIPA lysis buffer (Sigma-Aldrich) supplemented with protease and phosphatase inhibitor (ThermoFisher), incubated on ice for 10min, then shredded through QiaShredder columns (Qiagen) through centrifugation at 13,000RPM, 4°C for 2 min. Protein in the whole cell lysates was quantified using the Bradford method, normalized and prepared with 4X Laemmli Sample Buffer (Bio-Rad). Proteins were run on Mini-PROTEAN TGX Stain-Free Precast 4-20% Gels (Bio-Rad) and fast electrophoretic transferred to PVDF membrane (TransBlot Turbo, Bio-Rad). Membranes were blocked and probed in 5% BSA overnight at 4°C with primary antibodies as follows, β-actin (1:2000, CST#4970), c-MYC (1:500, Ab#32072), CNOT1 (1:1000, CST #44613), EGFR (1:1000, CST #2232), phospho-EGFR (Tyr1068) (1:1000, CST#2234), MCL-1 (1:1000, CST #94296), p300 (1:200, SC#32244), Rpb1CTD (1:1000, CST #2629), phospho-RPB1(Ser2) (1:1000, CST #13499), 4E-BP1(1:1000, CST #9644), phospho-4E-BP1 (Thr37/46) (1:1000, CST #2855), phospho-4E-BP1 (Ser65) (1:1000, CST #9451), phospho-4E-BP1 (Thr70) (1:1000, CST #13396), ubiquitin (1:1000, CST#3936), vinculin (1:2000, CST #13901). HRP-linked anti-Rabbit (CST #7074) and anti-Mouse (CST #7076) secondary antibodies were applied accordingly at a 1:5000 dilution in 5% milk in PBS-T at room temperature. Quantification of immunoblots was performed where indicated with ImageJ software (Rueden et al., 2017; Schneider et al., 2012). Background measurement was subtracted, and band intensity was normalized to loading control intensity.

### Immunoprecipitation

Cells were seeded in 15-cm plates, treated with DMSO or indicated drugs for 18h, to yield at least 1 mg of total protein for immunoprecipitation. All subsequent steps were performed on ice. At the time of harvest, cells were washed with PBS, pelleted (3000RPM, 4°C, 5min), resuspended and incubated on a rotator for 1h, 4°C in IP buffer (150 mM NaCl, 0.5% NP-40, 20 mM EDTA, 1 mM Dithiothreitol (DTT), 40 mM Tris HCl, pH7.4) supplemented with protease/ phosphatase inhibitor cocktail (ThermoFisher). After lysis was completed, lysates were clarified at 13000RPM, 4°C, 20min. Protein was quantified using the Bradford method and normalized to the lowest protein amount among the samples. Input controls were also saved and prepared with 4X Laemmli Sample Buffer (Bio-Rad) accordingly. 2-4 μg of primary antibodies (c-MYC, Ab#32072) or appropriate isotype control were added to respective samples and incubated overnight on a rotator at 4°C. 40 μL/sample of recombinant Protein-G-Sepharose-4B beads (ThermoFisher) was washed thrice with IP buffer and added to each sample for equilibration on a rotator for 4h, 4°C. At the end of the antibody-bead conjugation, immunoprecipitates were collected by centrifugation 3000RPM, 4°C, 5min. The bead pellets were washed for a total of 5 times. After the last wash, immunoprecipitated proteins were eluted with 4X Laemmli Sample Buffer, vortexed briefly and heated at 95°C, 5min. Samples were collected (13000RPM, 2min), subjected to SDS-PAGE and transferred to PVDF membrane as described above.

### RNA-sequencing sample preparation and analysis

RNA-sequencing was performed with ERCC spike-in normalization as previously described (Lovén et al., 2012). Briefly, CAL51 and MDA-MB-231 cells were seeded in 10-cm plates and incubated in media with DMSO or indicated drugs for 12h, in triplicates. Cell counts were determined using C-Chip disposable hemocytometers (Bulldog bio, DHC-N01) and equalized across all samples before lysis and RNA extraction. Total RNA from 1E6 cells per replicate was isolated using RNeasy96 kit (Qiagen). External RNA Controls Consortium (ERCC) ExFoldRNA Spike-in Control Mixes (Invitrogen #4456740) (4 μL/sample, diluted at 1:100, ERCC User Guide, Table 4) were added after cell lysis step. The extraction was then continued according to manufacturer’s instructions and eluted in 50 μL nuclease-free water. Total RNA was quantified using Qubit^TM^ RNA Broad Range Assay kit (Invitrogen) and analyzed on Agilent 4200 TapeStation for integrity. Samples with the RNA Integrity Number (RIN) above 9.0 and normalized to 500 ng total RNA for library preparation using TruSeq^®^ stranded mRNA sample prep kit (Illumina, #20020595). After library preparation, samples were quantified using Qubit^TM^ assay, checked fragment sizes on Agilent 4200 TapeStation, normalized and pooled. Libraries were sequenced on Illumina HiSeq 2000 sequencing system using 50-bp single-end reads at the Duke University Genome Sequencing Facility.

Sequences were processed using Trimmomatic v0.32 (Bolger et al., 2014) and reads that were 20nt or longer after trimming were filtered for further analysis. Reads were aligned using the alignment tool STAR v2.4.1a (Dobin et al., 2013) following the proposed 2-pass strategy to first identify a splice junction database to improve the overall mapping quality. Alignment was performed to GRCh38/hg38 of the human genome and transcriptome with ERCC synthetic spike-in RNA sequences (Annotations from product webpage manuals, https://assets.thermofisher.com/TFS-Assets/LSG/manuals/cms_095047.txt) appended for mapping. The TPM (transcripts per million) was computed for each mapped gene and synthetic spike-in RNA using RSEM v1.2.25 (Li and Dewey, 2011). Differential expression analysis was performed using DESeq2 v1.22.0 (Love et al., 2014) running on R (v3.5.1). Briefly, raw counts were imported and filtered to remove genes with low or no expression, that is, keeping genes having two or more counts per million (CPMs) in two or more samples. Filtered counts were then normalized with the DESeq function, using the counts for the ERCC spike-in probes to estimate the size factors. In order to find significant differentially expressed genes, the nbinomWaldTest was used to test the coefficients in the fitted Negative Binomial GLM using the previously calculated size factors and dispersion estimates. Genes having a Benjamini-Hochberg false discovery rate (FDR) less than 0.05 were considered significant (unless otherwise indicated). Differential gene expression was tested for all possible drug pairwise comparisons within each cell line, for example, single drug versus DMSO control, combination versus DMSO control, combination versus single drug and so on.

### Quantitative real-time PCR

Cell counts were determined and normalized across all samples before lysis. RNA was extracted using RNeasy Mini kit (Qiagen). After the cell lysis step, samples were spiked-in with the ERCC spike-in controls (2 μL/sample, diluted 1:100, Invitrogen #4456740), treated with on-column DNase digestion according to manufacturer’s specifications (Qiagen). RNA purity and concentration was measured by absorbance at 260nm (A_260_/A_280_). cDNAs were reverse-transcribed using SuperScript^TM^ VILO^TM^ cDNA Synthesis kit (Invitrogen) with 100 ng – 1 μg of RNA template as directed by the manufacturer’s protocol. qRT-PCRs were carried out in triplicates using TaqMan assay (Applied Biosystems) and CFX96 or CFX384 Touch Real-Time PCR Detection System according to manufacturers’ recommendations (Bio-Rad). Average cycle threshold (C_t_) values were calculated for each gene, and the maximum C_t_ value was set at 40 cycles. Average C_t_ values of technical replicates were normalized to the exogenous spike-in or reference gene, ERCC-00096 or GAPDH respectively, and relative gene expression was determined using the comparative ΔΔC_t_ method. Average and standard deviation were results of at least three independent experiments. Specific TaqMan gene expression assay IDs were as follows: ERCC-00096 (Ac03459927_a1), GAPDH (Hs02786624_g1), MYC (Hs99999003_m1), HSPA5 (Hs00607129_gH).

### Sucrose density gradient sedimentation of polysomes

CAL51 cells were seeded in 10-cm plates to reach 80 – 85% density at the point of harvest, treated with DMSO or indicated drugs for 12h. Before the time of harvest, gradients of 15% to 50% sucrose solutions (200 mM KCl, 15 mM MgCl_2_, 0.2 mM cycloheximide, 1 mM DTT, 10 U/mL RNaseOUT^TM^, 25 mM K-HEPES, pH7.4) were first prepared in centrifuge tubes. All subsequent steps were performed on ice. Untreated or treated cells were washed twice with ice-cold PBS and lysed on ice for 10min with 1 mL/ plate lysis buffer (200 mM KCl, 15 mM MgCl_2_, 1% NP-40, 0.5% sodium deoxycholate, 0.2 mM cycloheximide, 1 mM DTT, 40 U/mL RNaseOUT^TM^, 1X protease inhibitor, 25 mM K-HEPES, pH7.4). Lysates were clarified at 13000RPM, 4°C, 10min. Reserved supernatants were overlaid onto sucrose gradient. The samples were centrifuged in a swinging-bucket rotor SW41 Ti (Beckman) at 35,000RPM, 4°C, 3h with vacuum. Sample fractions were collected from the bottom of the tube with simultaneous absorbance A_260_ tracing (TracerDAQ^TM^) and subjected to RNA extraction followed by qRT-PCR analysis using TaqMan assays as listed above.

C_t_ values from qRT-PCR was analyzed to obtain relative distribution of respective mRNA as previously described (Pringle et al., 2019).

### [^35^S]methionine labeling

CAL51 cells were seeded in 6-well plates to reach 80-85% density at the point of labeling, treated with DMSO or indicated drugs. After 12h of vehicle or drug incubation, cells were washed twice with PBS and methionine-starved in serum-supplemented methionine-free media (Gibco) for 30min at 37°C. Cells were then labeled with 150 μCi/mL [^35^S]methionine (Perkin-Elmer) in methionine-free media for 45min. To stall translation and quench methionine incorporation, cells were washed twice with 100 μg/mL cycloheximide (Sigma-Aldrich) serum-free methionine-free media, incubating for 10min at 37°C for the second wash, followed by two washes with 100 μg/mL cycloheximide in PBS. Cells were then lysed with IP buffer on ice for 10min. Samples were pre-cleared by rotating with rabbit IgG isotype control (Ab #172730) and Protein-G-Sepharose-4B beads for 1h at 4°C. MYC immunoprecipitation were performed as described earlier. 2 μL of each samples were added to 3 mL of liquid scintillation liquid and [^35^S] radioactivity was measured and recorded. The remaining of the samples were subjected to SDS-PAGE and transferred to PVDF membrane as described above. Membranes were exposed on a phosphorimaging screen overnight and visualized on Amersham^TM^ Typhoon^TM^ NIR Biomolecular Imager (GE Healthcare).

### Cloning of CRISPR and shRNA constructs

CRISPR constructs were cloned using published methods (Shalem et al., 2014, Joung et al., 2017) using characterized sgRNAs from TKOv3 genome-wide library (Hart et al., 2017). Detailed cloning steps were as previously described (Lin et al., 2019). In brief, unique 20-mer sgRNA inserts targeting genes of interest were synthesized by CustomArray with flanking sequence adaptors. The synthetic oligo (diluted 1:100) was amplified using NEB Phusion Hotstart Flex PCR master mix and array primers. Amplified inserts were beads-clean-up at 1.8x (Axygen, AxyPrep^TM^ Mag PCR Clean-up kit) and eluted in molecular grade water. lentiCRISPRv2 (Addgene ID 52961) was digested by Esp3I (BsmBI) (ThermoFisher) and size selected by 1% agarose gel electrophoresis and extraction. Linearized lentiCRISPRv2 were then annealed with clean array amplified sgRNA oligos by Gibson assembly reaction. The reaction mixture was then transformed by chemical or electroporation method into Stbl3 (ThermoFisher) or E. cloni 10G (Lucigen) competent cells respectively. Transformed cells were recovered and spread on LB-ampicillin plates for overnight incubation. Single colonies were picked, cultured overnight in liquid LB and extracted using Plasmid miniprep kit (Qiagen). Plasmid DNA sequences were checked using Sanger sequencing (Eton Bioscience) for sgRNA inserts to confirm successful cloning. See Key Resources Table for sgRNA sequences.

Glycerol stocks for shRNA targeting genes of interest and bacterial stab cultures of plasmids were obtained from the Duke Functional Genomics Core Facility and Addgene Plasmid Repository respectively. Inoculants from glycerol stocks or stab culture were cultured overnight in liquid LB at 37°C and plasmids were extracted using Plasmid miniprep kit (Qiagen). See Key Resources Table for shRNA identity, TRC number and target sequences, and plasmid Addgene ID.

### Lentivirus production and transduction

Lentivirus production was adapted from Joung et al., 2017. HEK293FT cells were grown to ∼80% confluency in 10-cm or 6-well plates, for 10 mL or 2 mL final viral media harvest respectively and transfection reagents were scaled according to seeding area. For 10-cm plate, 3.5 – 4E6 cells were seeded and incubated for 24h (37°C, 5% CO2). Transfection reagents were prepared in Opti-MEM^TM^ reduced serum medium (Gibco) and performed using 94.2 μL Lipofectamine 2000 (ThermoFisher), 103.6 μL PLUS^TM^ reagent (ThermoFisher), 8.2 μg psPAX2, 5.4 μg pMD2.G and 10.7 μg construct DNA. The mixture was incubated at room temperature for 5min and gently added to the HEK293FT cells for 4h incubation (37°C, 5% CO2). The medium was then replaced with pre-warmed harvest media (DMEM 30% FBS). 48h after the start of the transfection, lentivirus supernatant was collected and syringed through a 0.45μm filter. Transductions were conducted directly at the time of lentivirus harvest or freshly thawed from frozen aliquots. 0.5-1 mL of virus media and polybrene (1 μg/mL) were added to cells seeded in 6-well plate in 1-1.5 mL of growth media. Cells were spinfected at 2250RPM, 1h, room temperature (25°C) and incubated overnight (37°C, 5% CO2). 24h post-transduction, cells were selected by puromycin (2 μg/mL) for 48h.

### Pooled genome-wide CRISPR positive selection screen and analysis

TKOv3 pooled library was obtained from Addgene (Addgene ID 90294) and amplified as previously described (Hart et al., 2017, Joung et al., 2017). Lentivirus production of TKOv3 library was scaled up and conducted as described above. CAL51 cells were seeded into 6-well plates at a density of 0.5E6 cells per well and transduced at a MOI less than 0.2. A total 60E6 cells were transduced in 24×6-well plates. 24h post-transduction, cells were selected by puromycin (2 μg/mL) for 48h. Puromycin-selected cells were collected and counted to confirm at least 100X library coverage. Transduced cells were propagated puromycin-containing media for a total of 7 days and split into vehicle (DMSO) and gefitinib (750 nM) + THZ531 (100 nM) combination treatment conditions in duplicates. The screen was conducted over a total of 3 weeks, for approximately 15 cell doublings. Cells were counted, passaged with replenished drug every 3 days. Each treatment condition and replicate was represented by a minimum of 10E6 cells to maintain at least 100X library coverage (>100 cells per unique sgRNA) during each split throughout the screen. A total of 12E6 cells were collected at 48h post-puromycin exposure, screen initiation (t_0_) and at every passage till screen termination (t_final_). DNA was extracted from cell pellets (DNeasy Blood & Tissue Kit, Qiagen) and stored at −80°C until completion of screens. Samples were further processed for sequencing as previously described (Shalem et al., 2014). Screen libraries were sequenced on Illumina NovaSeq 6000 sequencing system (50-bp, single-end reads) at the Duke University Genome Sequencing Facility to achieve 20 million reads total per sample (∼200 reads per guide).

Pooled samples were matched by barcoded reads and guide-level counts were computed using bcSeq (v1.12.0) Bioconductor package (Lin et al., 2018) in the R (v3.5.1) programming environment. As the screen was designed for positive selection, resistance to gefitinib+THZ531 combination was determined by evaluating differential guide compositions between vehicle control (DMSO) and combo-treated (GT) cell populations at t_final_. Cells that survived the GT combo were enriched with guides targeting genes that we coined ‘resistor’ genes and are required for the drug synergistic activities. Differential analysis was carried out using the DESeq2 (v1.22.0) Bioconductor package in the R (v3.5.1) programming environment. Out of the 18,053 genes in the TKOv3 library, 29 genes (0.16%) were excluded due to low counts. Enrichment effects in combo-treated arm were expressed as log_2_(fold-change) for GT versus DMSO (vehicle-control as the denominator).

### Quantification and Statistical Analysis

Statistical analyses were performed in Prism9 (GraphPad) software or R (v3.5.1) (https://www.r-project.org/). All results are shown as mean ± standard deviation. P-values were determined using unpaired, two-tailed Student’s t-tests and considered significant at threshold of <0.05, unless otherwise stated.

### Data and Software availability

All raw data from the study are either supplied in the Supplemental Information or will be made available upon request. RNA-seq data for CAL51 and MDA-MB-231 treated with the specified drug conditions are deposited onto the Gene Expression Omnibus (GEO accession numbers supplied when available). RNA-seq counts table after renormalization with synthetic ERCC spike-in-RNA is available in Supplemental Table S1A and S1B. Raw counts table for the TKOv3 positive selection screen of CAL51 cells treated with DMSO or gefitinib+THZ531 combination is available in Supplemental Table 2. Analyzed screen data is available in Supplemental Table 3.

